# Proton motive force mediated efflux mismatch drives gentamycin-novobiocin collateral sensitivity in *Pseudomonas aeruginosa*

**DOI:** 10.64898/2026.05.01.722319

**Authors:** Hafij Al Mahmud, MD Siddiqur Rahman, Erika Ortiz, Alex Luecke, Amanda M.V. Brown, Catherine Wakeman

## Abstract

Resistance to a particular antibiotic can make bacteria sensitive to others, a phenomenon known as collateral sensitivity (CS). This study explored potential CS in clinical and experimentally evolved drug-resistant *Pseudomonas aeruginosa* (PA) and investigated underlying mechanisms. Whole-genome sequencing and RNA-seq were analyzed to identify genetic and transcriptional correlations. *In vitro* efficacies were assessed with co-and sequential-exposure regimens. Multiple CF isolates and experimentally evolved gentamycin (GEN) resistant strains consistently exhibited strong CS to novobiocin (NOV). Comparative genomics revealed *pmrB* gain-of-function mutations, which was further supported by transcriptomic signatures of pmrAB activation. Transcriptomic data suggests potential outer-membrane remodeling characterized by polyamine accumulation and compromised porin channel expression. Additionally, the reduction in proton motive force (PMF) further explains the possible mechanism underlying GEN resistance. As NOV efflux is PMF-dependent, this energetic deficit created a PMF-efflux mismatch, leading to hypersensitivity to NOV. Notably, sequential GEN→NOV treatment effectively restricted the emergence of GEN resistant subpopulations. Overall, our data suggest GEN resistance in PA may arises through envelope remodeling and reduced PMF, which impairs efflux pumps and creates hypersensitivity to NOV. Exploiting this PMF-efflux mismatch with sequential treatment effectively restricted the emergence of GEN resistance.

## Introduction

Combating emerging antibiotic resistance in clinical settings is a major challenge worldwide. In recent decades, the excessive utilization and inappropriate application of antibiotics have emerged as primary factors contributing to the rise of antibiotic resistance in bacteria (1, 2). For example, the prolonged administration of antimicrobial treatments prescribed for managing chronic infections like cystic fibrosis (CF) infection demands continual antibiotic exposure, that may amplify the evolution of multidrug or extensive drug resistance (3). Extensive utilization of antibiotics in agriculture, livestock farming, and animal husbandry practices are also significant driving factors (4). Additionally, the limited number of new antibiotic discoveries necessitates novel strategies to tackle bacterial infections.

Among the bacteria causing significant concern, *Pseudomonas aeruginosa* (PA) stands out as a notable pathogen responsible for a variety of life-threatening acute and chronic infections (5, 6). This bacteria acts as an opportunistic pathogen, causing nosocomial infections and persistent infections in people with diseases such as chronic obstructive lung disease or CF infection (7, 8). PA demonstrates remarkable antibiotic resistance through a combination of inherent and mutation-induced mechanisms, leading to multidrug or pan-drug resistance (9, 10). Multidrug efflux pumps play a major role as the antimicrobial-resistant determinants in microorganisms, including PA. They facilitate intrinsic, acquired, and transient antibiotic resistance by actively expelling various antimicrobial agents from bacterial cells (11–13).

The rapid emergence of antimicrobial resistance and the limited discovery of new antimicrobial agents necessitate the design of rational treatments from existing antimicrobials. To this end, a potential therapeutic approach involves antibiotic cycling, leveraging a phenomenon known as collateral sensitivity (CS) (3). CS is a phenomenon where resistance to one treatment leads to enhanced sensitivity to another, offering new possibilities to tackle the emergence of drug resistance (14, 15). Even though CS was first explained in the 1950s, it has gained significant momentum in recent days (16). For example, the scientific community is focused on exploiting CS to combat the emergence of antibiotic resistance by identifying new vulnerabilities in drug-resistant pathogens, including *Escherichia coli* (14, 17–19), *Staphylococcus aureus* (20), PA (19, 21), *Salmonella typhimurium* (22), *Burkholderia multivorans* (3), *Klebsiella pneumoniae* (23, 24), *Salmonella enterica* (25), *Acinetobacter baumannii* (26, 27), etc. This phenomenon is not restricted to bacterial cells. In fact, cancer cells have also shown that drug resistance evolution can alter susceptibility to unrelated drugs, either sensitizing or causing resistance (28). Recent studies have shown that phage–antibiotic resistance trade-offs can restore antibiotic sensitivity and extend the usefulness of existing drugs (29–31).

Utilizing CS in treatment strategies could be a useful method to prevent or even reverse the development of resistance (24). In this regard, antibiotic combination therapy based on CS could be a promising approach through which we can achieve extended antibiotic sensitivity and reduce the emergence of drug resistance. The efficacy of CS-based treatments can be influenced by several factors including genetic makeup of resistant mutants, molecular structure of antibiotics, environmental conditions, etc (15). Genetic background of the resistant mutant is crucial, for example, PA populations adapted to the same drug may acquire distinct resistance mutations (32), leading to different CS patterns due to variation in associated fitness trade-offs (33–36). The potential of CS-based therapies is considerable, offering a strategy to restrict the emergence of resistance and reduce infection relapse. In this study, we examined antibiotic-resistant PA clinical isolates from CF patients and identified a unidirectional CS between gentamicin (GEN) and novobiocin (NOV). We further investigated the underlying mechanisms of this resistance trade-off and proposed a sequential combination therapy to limit the reemergence of GEN resistance.

## Methods and Materials

### Evaluating antibiotic sensitivity via disc diffusion assay

The antibiotic susceptibility of PA strains was assessed using the disc diffusion method. Briefly, PA was cultured on lysogeny broth (LB) in a shaker for 16-20 hours at 37°C. Subsequently, bacterial cultures were normalized to an optical density of 0.1 at 600nm. Sterile cotton buds were saturated with the cell culture. After removing excess media, the cotton buds were used to evenly distribute the bacterial culture onto lysogeny agar (LA) plates. Antibiotic discs were carefully placed onto the plates, and zones of inhibition were observed following 24 and 48 hours of incubation at 37°C. The diameter of the zones of inhibition was measured in millimeters to evaluate the susceptibility of PA strains to the antibiotics tested.

### Generating antibiotic-resistant bacteria through experimental evolution

Antibiotic-resistant bacteria were generated through experimental evolution in the laboratory following a published protocol with slight modification (37). The overnight culture of *Pseudomonas aeruginosa* UCBPP-PA14 strain was diluted 100 times in 5 ml fresh LB medium in 15 ml tubes preoccupied with/without antibiotics. The culture was subsequently diluted every day and incubated at 37°C. The concentration of each antibiotic increased over time from suboptimal concentration to a very high concentration. Three consecutive cultures were carried out for a particular antibiotic concentration. Following the liquid culture, antibiotic-resistant bacteria were isolated from respective antibiotic-containing agar plates for further experimentation.

### Evaluating antibiotic sensitivity using broth microdilution assay

Antibiotic sensitivity was evaluated following a previously published protocol (38). Overnight PA cultures were adjusted to an optical density of 1.0 at 600nm. Finally, cells were diluted in LB to get a final cell density of around 5X10^5^ cells per ml, and 200 µl of the diluted cells were inoculated into 96-well plates. Various concentrations of antibiotics (GEN; max-100 µg/ml, NOV; max-1200 µg/ml) were added to the wells, and appropriate controls were set up. After 24 hours of incubation at 37°C, resazurin (Sigma Aldrich, USA), a fluorogenic dye, was added to each well and further incubated for 2 hours. Following incubation, the relative fluorescent units were estimated by using a multiple reader (Ex; 540, Em; 590). Following 3 hours of additional incubation, minimum inhibitory concentrations (MICs) were determined based on the color change, where the blue color depicts no growth, whereas the pink color depicts viable cells.

### Genomic DNA extraction, sequencing and comparative genomics

Parental strain PA14 and GEN resistant (GENr) bacteria were grown on LB and genomic DNA were extracted using DNeasy Blood and Tissue Kits (Qiagen, Germany) and finally sent for Illumina sequencing. Once the raw sequence data were received, genome comparison was performed following a published article (37). Briefly, parental and evolved genome sequences were filtered and trimmed using Trimmomatic software v0.38 with the following selected criteria: LEADING:20 TRAILING:20 SLIDINGWINDOW:4:20 MINLEN:100. Next, the variant calling was performed using the breseq software v0.38.2 (39) using the default parameters. The complete genome of *Pseudomonas aeruginosa* UCBPP-PA14 (NC_008463.1) was used as reference. Mutations present in the parental strain were subtracted from evolved strains using gdtools, and remaining variants were compared across GENr isolates.

### Efflux pump inhibition assay

Bacterial cultures were prepared to the desired inoculum density in phosphate buffered saline (PBS). The efflux pump inhibitor carbonyl cyanide m-chlorophenyl hydrazone (CCCP) was freshly prepared at 200 µM. Various concentrations of CCCP (12.5, 25 and 50 µM) were tested along with PBS control. Ethidium bromide (EtBr) was then added to all samples at a final concentration of 128 µg/ml. Finally, bacterial suspensions were added to each reaction mixture, resulting in a uniform final volume across all conditions.

### Evaluating intracellular NOV concentrations

Overnight cultures of PA14 and GENr strains were diluted 1:10 into fresh LB and incubated for 1.5 hrs at 37 °C with shaking. Cells were harvested, washed, and adjusted to an OD_600_ of 1 in 1XPBS. Aliquots of 50 µl were transferred to microcentrifuge tubes, and Novo-TRX (Texas Red-labeled novobiocin; BPS Bioscience, CA, USA) was added to a final concentration of 50 nM. Samples were incubated at 37 °C, 120 rpm for the desired time points, pelleted by centrifugation (16,000 × g, 1 min), washed once with 1XPBS, and resuspended in 50 µl 1XPBS. Fluorescence was measured at Ex 584 nm and Em 625 nm.

### RNA extraction and library preparation

*Test condition:* PA overnight cultures were normalized to an OD_600_ of 0.8, and 20 ml aliquots were supplemented with GEN (15.6 µg/ml), NOV (2400 µg/ml), along with no-drug control. Cultures were incubated for 1 h at 37 °C in a shaking incubator. Following incubation, cells were harvested by centrifugation at 5000 rpm for 5 min, and supernatants were discarded.

### RNA extraction

Bacterial pellets were lysed by lysozyme and further treated with proteinase K for 15 min at room temperature. Total RNA was extracted using the RNeasy Mini Kit (Qiagen) according to the manufacturer’s instructions. Purified RNA samples were quantified and the quality was checked using the Agilent TapeStation system.

### Library preparation and sequencing

Ribosomal RNA was removed using the RiboMinus Bacteria 2.0 Transcriptome Isolation Kit (Invitrogen) following the manufacturer’s protocol. RNA-seq libraries were then generated from rRNA-depleted RNA using the NEBNext Ultra II RNA Library Prep Kit for Illumina (New England BioLabs), according to the manufacturer’s protocol (Section 4: rRNA-depleted RNA). Library preparation steps included RNA fragmentation and priming, first- and second-strand cDNA synthesis, purification of double-stranded cDNA with Mag-Bind® RxnPure Plus (Omega BioTek, USA) Sample Purification Beads, end repair, adaptor ligation, and PCR enrichment. Libraries were purified at each stage using Mag-Bind® RxnPure Plus Sample Purification Beads, and quality was assessed on an Agilent TapeStation. Final libraries were submitted to Genewiz for sequencing on the Illumina HiSeq platform.

### RNA-seq analysis

RNA-seq datasets were processed using Rockhopper, which performed reference-guided transcript assembly with *P. aeruginosa* UCBPP-PA14 as the reference genome (40). Raw gene-level counts were used for differential expression analysis in DESeq2. Genes with an adjusted p-value (Benjamini–Hochberg) ≤0.05 and a fold change ≥2.0 were considered significantly differentially expressed. The Venn diagrams were generated using the online tool Venny 2.0 (https://bioinfogp.cnb.csic.es/tools/venny/) to visualize the overlap in gene expression in different conditions.

### Effect of NOV and GEN on PA14 under anaerobic conditions

PA14 were grown aerobically in LB at 37°C and then diluted to reach an OD_600_ of 0.3. The effect of GEN and NOV on PA14 under anaerobic conditions was evaluated using two different test conditions. The first condition included LB supplemented with 1% potassium nitrate (41) that can be used as an electron acceptor in the absence of oxygen under anaerobic conditions; therefore, the bacteria experienced active proton motive force (PMF). The second condition contained LB supplemented with 100 mM potassium phosphate and 40 mM pyruvate, in which bacteria underwent anaerobic fermentation (42) and experienced compromised PMF. In both conditions, the maximum concentrations tested for NOV and GEN were 1200 and 100 µg/ml, respectively. Bacterial growth was then assessed by enumerating colony-forming units (CFUs) per ml.

### Antibiotic combination efficacy

#### Co-treatment

Overnight PA14 cultures were adjusted to an OD_600_ of 1.0, then diluted 1:100 into 3 ml LB with/without antibiotics. Three conditions were tested: no-drug control, GEN alone, and GEN combined with NOV (0.5X; 200 µg/ml). GEN concentrations gradually increased from 0.5X to 1X and 5X MIC (1X MIC = 1.56 µg/ml), each maintained for three consecutive passages. Following co-treatment, cultures were serially diluted in 1XPBS and spoted on LA and LA+20X GEN plates. CFUs were enumerated after 24 hrs incubation at 37 °C.

#### Sequential

Overnight PA14 cultures were adjusted to an OD_600_ of 1.0, then diluted 1:100 into 200 µl LB in 96-well plates with/without GEN. Both GEN-adapted populations (GENr) and no-drug controls were tested in triplicate. GEN concentrations were progressively increased from 0.5X to 20X MIC. Each concentration was maintained for three passages, except 20X MIC (one passage). Following serial dilution with 1XPBS, cultures were spot plated on LA, LA+1X NOV, LA+20X GEN, and LA+1X NOV+ 20X GEN plates. CFUs were enumerated after 24 hrs incubation at 37 °C.

## Results and Discussion

The rapid emergence of antibiotic resistance is one of the major challenges in treating chronic infection, which causes significant mortality across the world and costs a substantial amount of money (43). In addition, the trivial number of new antibiotics launched in the market for the past few decades necessitates finding new ways to utilize existing antibiotics to tackle the antibiotic resistance threat (44). To this end, we wanted to identify evolved vulnerabilities in drug-resistant pathogens to combat infection.

### Identifying vulnerabilities in CF clinical isolates of PA

We tested the antibiotic susceptibility of cystic fibrosis clinical isolates of PA (CFPA) together with wild-type lab strain PA14 against different antibiotics. Among them, we found an interesting CS between GEN and NOV in CFPA (data not shown). To further validate this CS, we performed disc diffusion assay. In this susceptibility test, unlike PA14, most of the CFPA isolates including CF666 and CF2577 showed increased resistance towards GEN and susceptibility to NOV (Fig 1A). Clearing zone diameters (millimeters) were measured at 24 and 48 hours of incubation and compared to PA14, with changes in zone values visualized as a heatmap (Fig 1B). While a few strains, such as CF2540, showed sensitivity pattern similar to PA14 (sensitive to GEN and resistance to NOV), most CFPA isolates exhibited interesting CS characteristics (resistant to GEN but susceptible to NOV) (Fig 1B).

**Figure 1.**
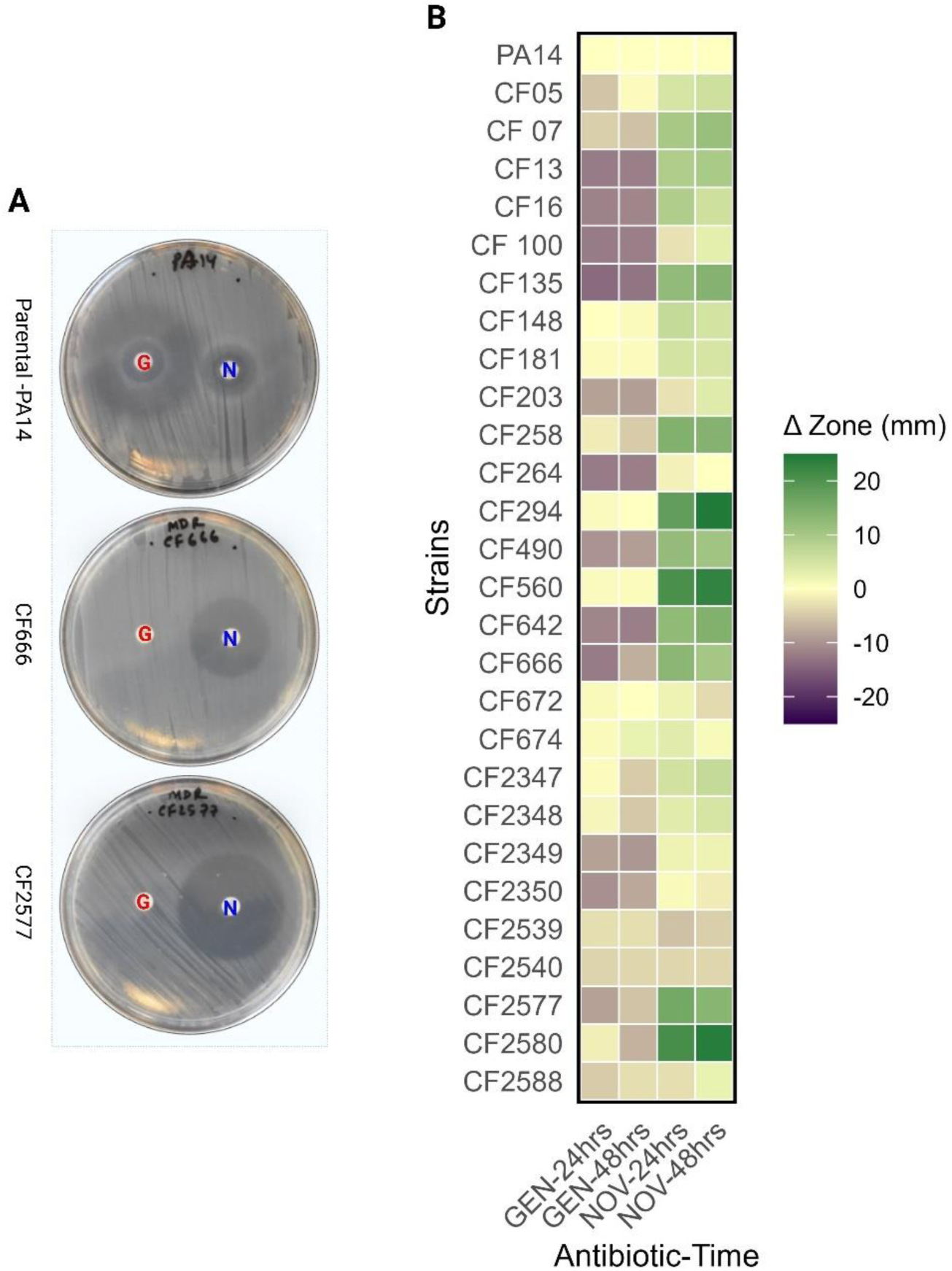
Identification of collateral sensitivity to gentamicin and novobiocin in evolved CF isolates. (A) Zones of inhibition for gentamicin (GEN) and novobiocin (NOV) against parental PA14 and CF isolates using the disc diffusion method. (B) Difference in inhibition zone diameters (mm) compared to PA14. In heatmap, green indicates increased sensitivity, red indicates increased resistance. Experiments were performed in triplicate on three independent days. Abbreviations: PA, *P. aeruginosa*; NOV, novobiocin; GEN, gentamicin.

To further validate the observed CS, we generated GENr cells through experimental evolution (Fig 2A). The broth microdilution method was used to determine the NOV and GEN MIC against PA14, GENr, and CFPA isolates (Fig 2B). MICs were normalized to PA14, and fold changes were visualized as a heatmap, where purple indicates increasing resistance, while green indicates increasing sensitivity. All four tested GENr strains exhibited resistance to GEN and increased sensitivity to NOV compared to the parental PA14 strain. This supports the earlier disc diffusion results, indicating that evolved resistance to GEN may enhance susceptibility to NOV. Among the 27 tested CFPA isolates, 8 isolates (e.g., CF13, 16, 100, 264, 490, 560, 666, and 2577) exhibited strong resistance to GEN and increased sensitivity to NOV. Additionally, 13 isolates showed elevated GEN resistance, and of those, 8 also displayed increased NOV sensitivity, also demonstrating a collateral sensitivity pattern. A few strains, including CF148, 181, 294, 672 and 674, remained sensitive to both antibiotics (Fig 2B). To complement the MIC data, we evaluated the relative fitness of PA14, GENr strains, and a representative clinical isolate CF666 using fluorescence-based assay, where higher fluorescence indicates greater cell viability (45). Overall, all GENr strains and CF666 exhibited significantly (*p<*0.05-0.0001) increased resistance to GEN compared to PA14 in a concentration-dependent manner (Fig 2C). In contrast, both GENr strains and CF666 showed significantly (*p<*0.05-0.0005) heightened susceptibility to NOV in a concentration-dependent manner (Fig 2D). These complementary trends, observed across MIC, disc diffusion, and fluorescence-based viability assays, consistently demonstrate a CS relationship between GEN and NOV. These results suggest that evolved resistance to GEN may enhance susceptibility to NOV in PA.

**Figure 2.**
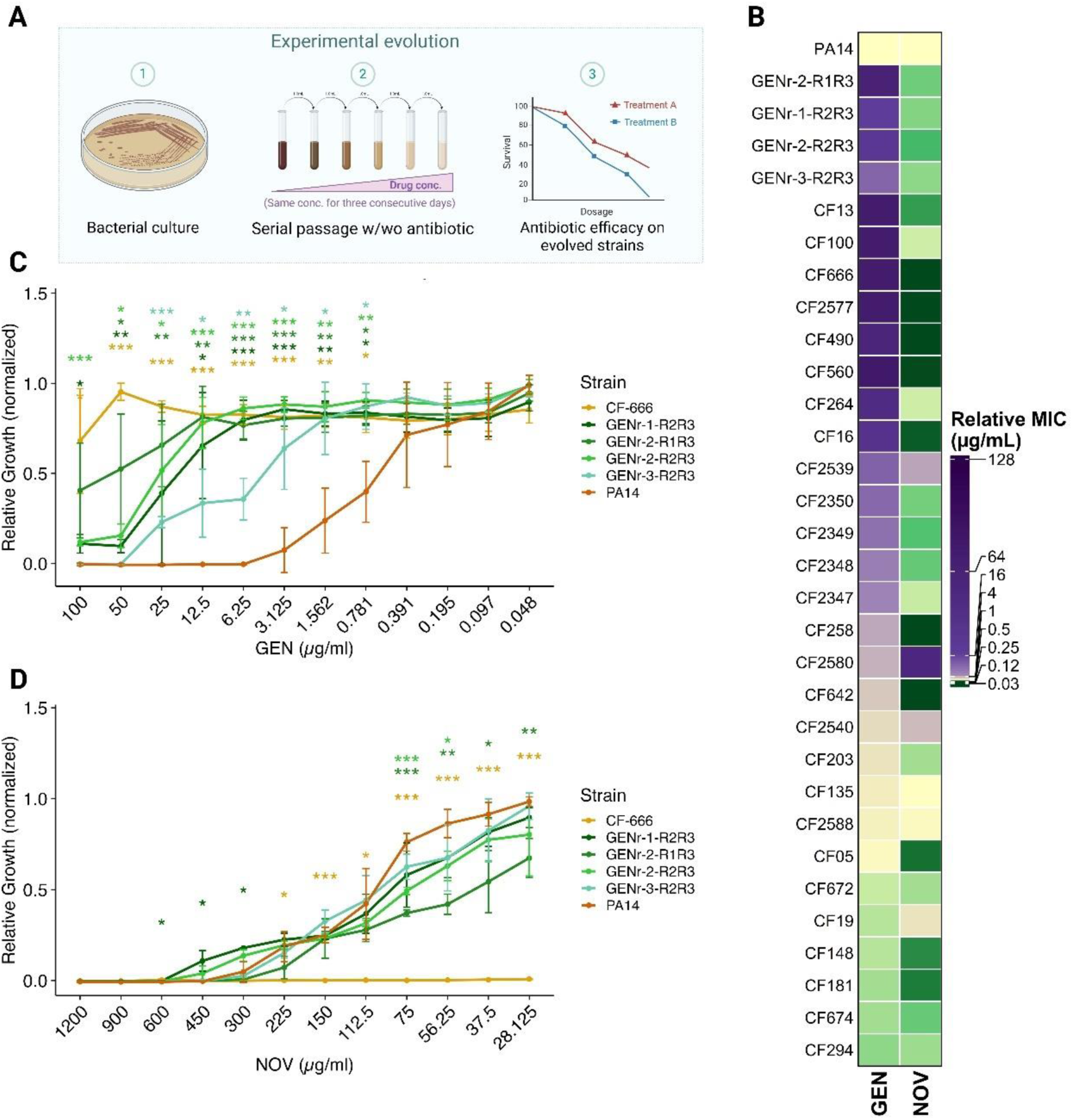
Validating collateral antibiotic sensitivity in experimentally evolved GENr strains through broth microdilution method. (A) Workflow for generating experimentally evolved gentamicin-resistant *P. aeruginosa* strains via serial passage. (B) Fold changes in MIC values for each strain relative to PA14. Green indicates increasing sensitivity, while purple indicates increasing resistance. Relative fitness of each strain following (C) GEN and (D) NOV treatment, compared to no-drug controls. All experiments were performed in triplicate on three days. Abbreviations: PA, *P. aeruginosa*; NOV, novobiocin; GEN, gentamicin. Besides, ‘*’ denotes *p* < 0.05, ‘**’ denotes *p* < 0.005, ‘***’ denotes *p* < 0.0005, and ‘****’ denotes *p* < 0.0001, as depicted by a two-tailed unpaired Student’s t-test.

After identifying GEN–NOV CS in PA, we tested whether this pattern extends to other pathogens by generating GENr *Acinetobacter baumannii*, *E. coli*, and *S. aureus* through experimental evolution. A similar GEN–NOV CS was observed in *S. aureus* but not in the other species (S1 Fig), indicating that this phenomenon is not restricted to PA or Gram-negative bacteria. Further studies are needed to clarify the mechanism enabling CS in *S. aureus*. Even though, CS was first described in *E. coli* in 1950 (16), but it has received increased attention and appreciation in recent years due to its potential to design more promising therapies against life-threatening infections. The CS between different antibiotics has been studied by many scientists across the globe on different clinically important pathogens such as *S. aureus* (20), PA (19), *S. typhimurium* (22), *B. multivorans* (3), and so on. For example, increased resistance to ciprofloxacin in PA may make them vulnerable to other antibiotics such as tobramycin (9, 46, 47).

### Identifying genes responsible for CS between GEN and NOV

To identify genes responsible for the observed CS in PA, we performed whole-genome sequencing of four GENr strains, alongside a no-drug experimental evolution control strain (EE-PA14) and the parental PA14 strain. The genomic alterations of the GENr strains were then compared to both the control and parental strains (Fig 3A). From this comparative genomic analysis, we identified four potential genes, including *EF-tu* (elongation factor tu), *pmrB* (two-component system sensor histidine kinase PmrB), 30s ribosomal protein S6/23S rRNA (guanosine(2251)2’-O)-methyltransferase *RimB* and a hypothetical protein (Table 1). A point mutation (A→C) has been detected in position 5,637,077 on the genome (particularly on *pmrB*) of all the GENr strains compared to the parental genome (Fig 3B). Besides, the amino acid H (histidine) has been changed to P (proline) on the PmrB amino acid sequence for GENr strain compared to the parental strain (Fig 3B). Among the potential gene targets, we found *pmrB* very interesting and wanted to explore further. Earlier studies reported an increased sensitivity to penicillin by GEN-adapted CFPAs because of the functional mutation on the *pmrB* gene (48, 49). Besides, it has been found that *pmrB*-mediated aminoglycoside resistance in *E. coli* can be regulated by the regulatory systems PhoP-PhoQ and PmrA-PmrB, which may make them sensitive to b-lactams antibiotics (50, 51). These regulatory systems can reduce the negative charges in the bacterial membrane by altering the lipid A profile in the bacterial outer membrane (50, 51). Other studies have also identified that functional mutations in *prmAB* genes may reduce the transmembrane potential of bacteria (52). The uptake of aminoglycoside antibiotics such as gentamicin depends on the functional PMF or transmembrane potential in bacteria (24, 53–56). Based on our comparative genomics analysis and prior studies, we hypothesize that a functional mutation in *pmrB* may reduce the PMF in PA. This reduction in PMF may impair the uptake of GEN while simultaneously limiting the PMF mediated efflux of NOV, thereby conferring resistance to GEN and increased sensitivity to NOV.

**Fig 3.**
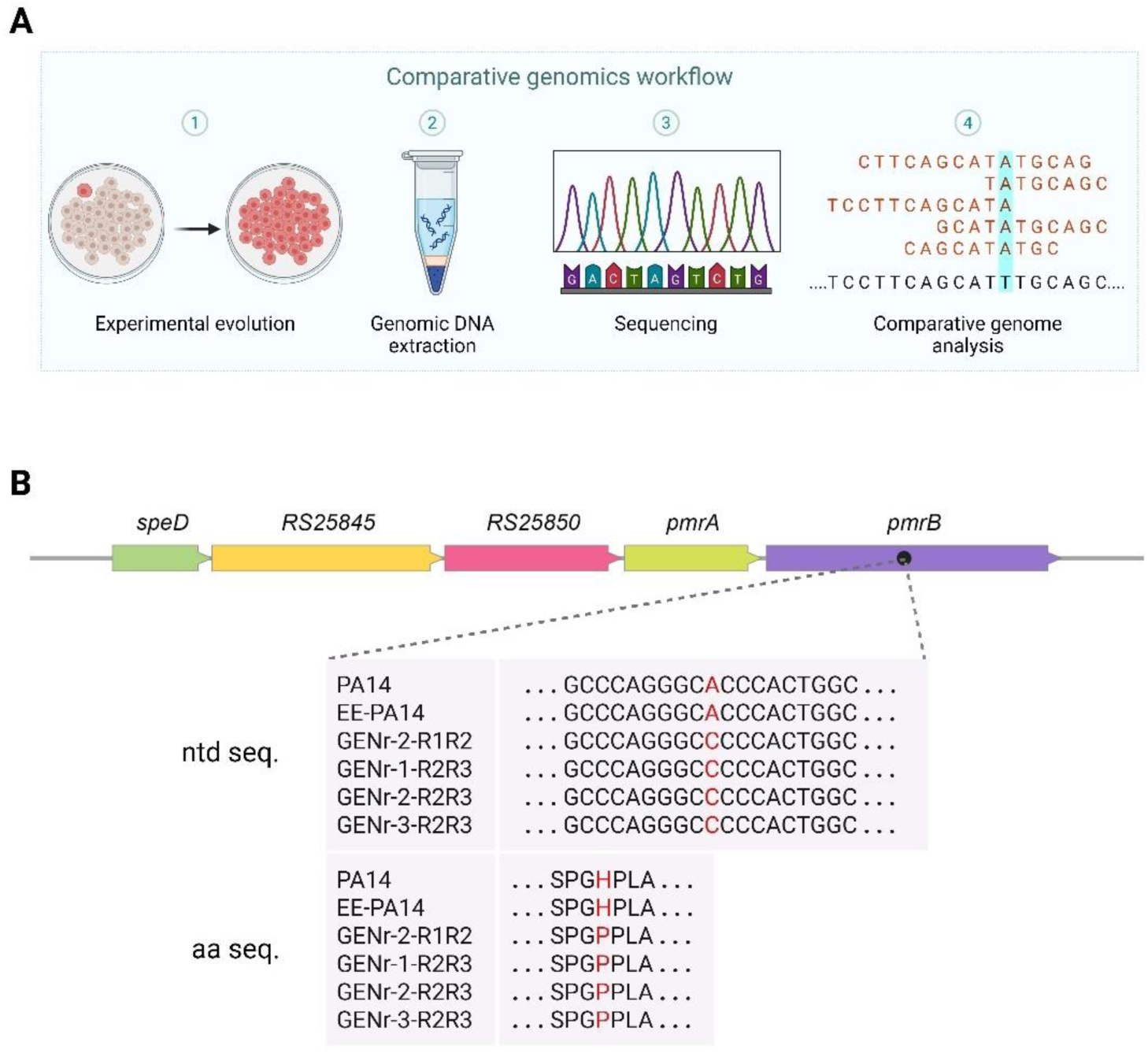
Comparative genomics to identify mutations associated with gentamicin resistance. (A) Workflow used to compare the genomes of GENr strains, the no-drug control (EE-PA14), and the parental PA14 strain. (B) Comparative genomics identified single nucleotide polymorphisms (SNPs) unique to gentamicin-adapted strains, distinguishing them from the parental and control strains.

**Table 1.** List of probable genes responsible for GEN-NOV CS predicted by comparative genomics analysis by breseq bioinformatic tools.

| Position | Mutation | Annotation | Gene | Description |
| --- | --- | --- | --- | --- |
| 741,564 | A→G | H85R<br>(CAC→CGC) | *tuf* | elongation factor Tu |
| 741,651 | C→T | P114L<br>(CCG→CTG) | *tuf* | elongation factor Tu |
| 3,843,716 | C→G | A124A<br>(GCG→GCC) | *PA14_RS17530* | hypothetical protein |
| 5,637,077 | A→C | T132P<br>(ACC→CCC) | *pmrB* | two-component system sensor histidine kinase PmrB |
| 5,809,296 | Δ1 bp | intergenic<br>(-182/+37) | *rpsF / rlmB* | 30S ribosomal protein S6/23S rRNA (guanosine(2251)-2'-O)-methyltransferase RlmB |

### Proton motive force drives GEN-NOV collateral sensitivity

To validate the role of PMF in the observed GEN-NOV CS in PA, we measured transmembrane potential by monitoring EtBr sequestration in GENr strains and PA14 over 60 minutes at 1-minute intervals, with or without CCCP treatment. CCCP, a known PMF uncoupler, was used at concentrations ranging from 12.5 to 50 µg/ml. Following 30 minutes of incubation, in the absence of CCCP, GENr strains exhibited higher EtBr accumulation than PA14. This result is consistent with our hypothesis that reduced PMF in GENr cells limits EtBr efflux (Fig 4A). Interestingly, without CCCP, PA14 showed higher initial EtBr uptake, likely due to its higher metabolic activity compared to GENr cells. Upon CCCP treatment, PA14 displayed a concentration-dependent increase in EtBr accumulation, surpassing that of GENr strains, further supporting the notion of diminished PMF in GEN-adapted bacteria (Fig 4A). Earlier studies also reported that, functional mutations in *pmrAB* can reduce the transmembrane potential of bacteria (52).

**Fig 4.**
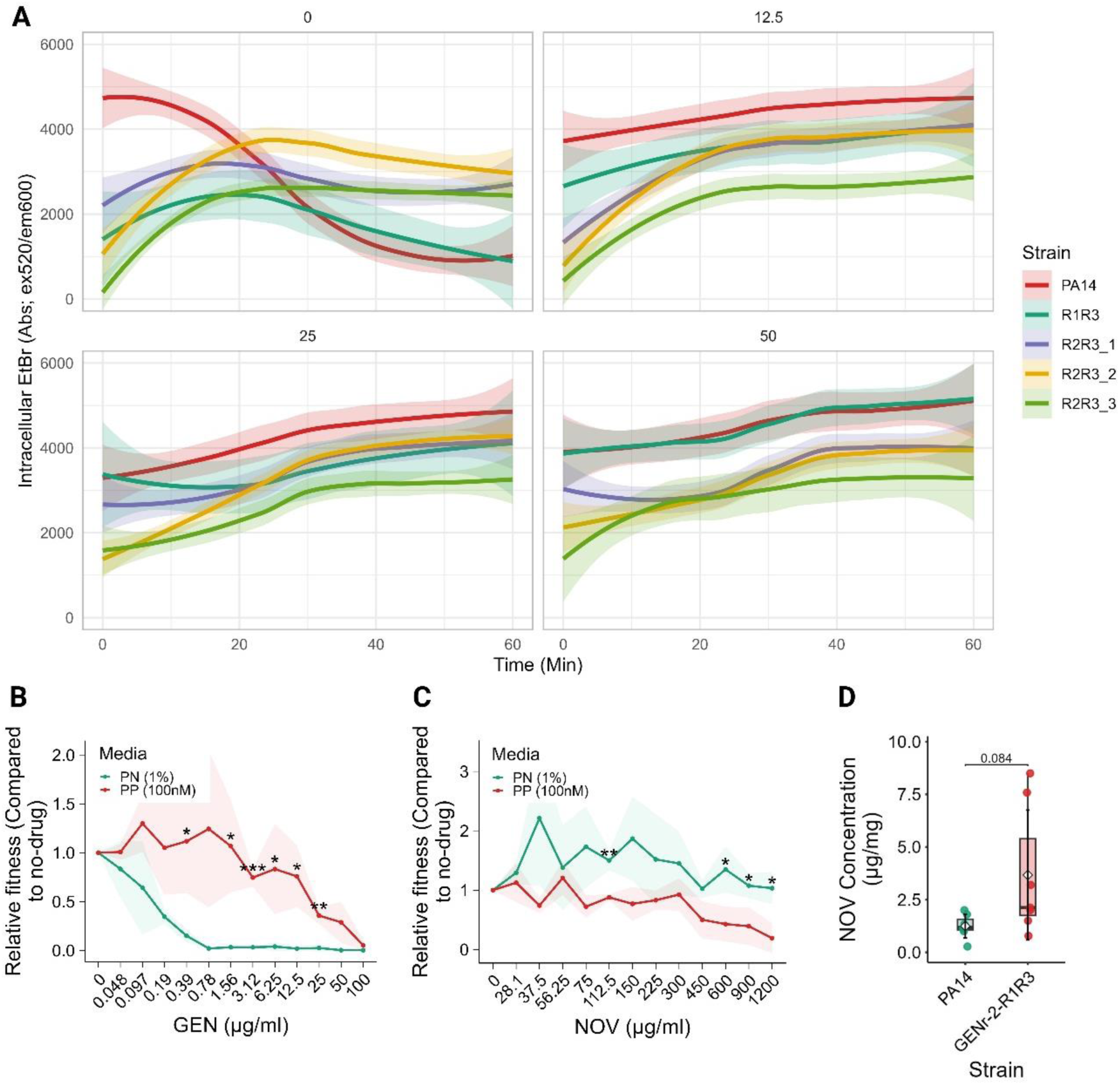
Role of proton motive force on GEN resistance and NOV sensitivity. (A) Accumulation of intracellular EtBr (ethidium bromide) by PA14 and GENr strains in presence or absence of CCCP (carbonyl cyanide m-chlorophenyl hydrazone). Inhibition of efflux pump was assessed using 128 µg/ml EtBr with 0-50 µg/ml CCCP. The sensitivity of PA14 to GEN (B) and NOV (C) in anaerobic respiration and fermentation conditions. Media supplemented with 1% potassium nitrate facilitates anaerobic respiration and 100 mM potassium phosphate and 40 mM pyruvate facilitates anaerobic fermentation. Data was obtained in triplicates on three different days. The error bar symbolizes the SD of the data. GEN; gentamicin, NOV; novobiocin, PN; potassium nitrate, and PP; potassium phosphate. Besides, ‘*’ designates *p* < 0.05, ‘**’ designates *p* < 0.005, and ‘***’ designates *p* < 0.0005, as depicted by a two-tailed unpaired Student’s t-test.

To further validate the role of PMF in GEN-NOV CS, we conducted experiments under anaerobic conditions considering two distinct metabolic states. In one condition, anaerobic fermentation was achieved by supplementing potassium phosphate-buffered media with pyruvate (42). In the other, anaerobic respiration was achieved by adding potassium nitrate, which serves as an alternative terminal electron acceptor in the absence of oxygen (57). In general, PMF is found to be reduced during anaerobic fermentation compared to aerobic or anaerobic respiration [42, 61]. We assessed the susceptibility of the PA14 strain to GEN and NOV under both conditions. Compared to anaerobic respiration, PA14 exhibited increased resistance to GEN under anaerobic fermentation, where PMF is compromised (Fig 4B). Conversely, PA14 showed increased sensitivity to NOV under anaerobic fermentation, further supporting the link between reduced PMF and GEN-NOV CS (Fig 4C). Additionally, following NOV treatment in aerobic culture, we observed an increased concentration of NOV in GENr bacteria compared to parental PA14 strain (Fig 4D).

Together, these results demonstrate that reduced PMF plays a key role in the CS between GEN and NOV in PA. GENr strains exhibit diminished PMF, leading to decreased GEN uptake and impaired NOV efflux, thereby conferring resistance to GEN and increased sensitivity to NOV.

### Transcriptomic analysis reveals energy mismatch underlining NOV-GEN collateral sensitivity

Venn diagrams illustrate the distribution of differentially expressed genes (DEGs) in GEN-adapted PA under no-drug, GEN, and NOV treatments (Figs 5A, S2). The majority of DEGs were found to be condition specific. For example, around 61.1% of downregulated genes unique to GEN treatment, 23.7% unique to NOV treatment, and 6.4% unique to the no-drug control (**Supplemental file 1**). Similarly, 60.1% of upregulated genes were unique to GEN treatment, 27.6% to NOV treatment, and 2.2% to the no-drug control. This implies that the observed CS does not solely result from genomic mutations, instead drug-specific reprogramming of gene expression might have significant role. Overlap between conditions was modest, for example, only 6.2% of downregulated and 3.4% of upregulated genes were shared between GEN and NOV treatment, while just 0.5% of downregulated and 4.2% of upregulated genes were common to all three conditions (Fig 5A). The small set of common genes likely represent baseline adaptations of GENr PA regardless of treatment. Overall, these patterns indicate that while a small set of baseline responses is maintained, the transcriptional programs of GENr strains are predominantly shaped by the specific treatment, with only limited overlap across treatment conditions.

**Fig 5.**
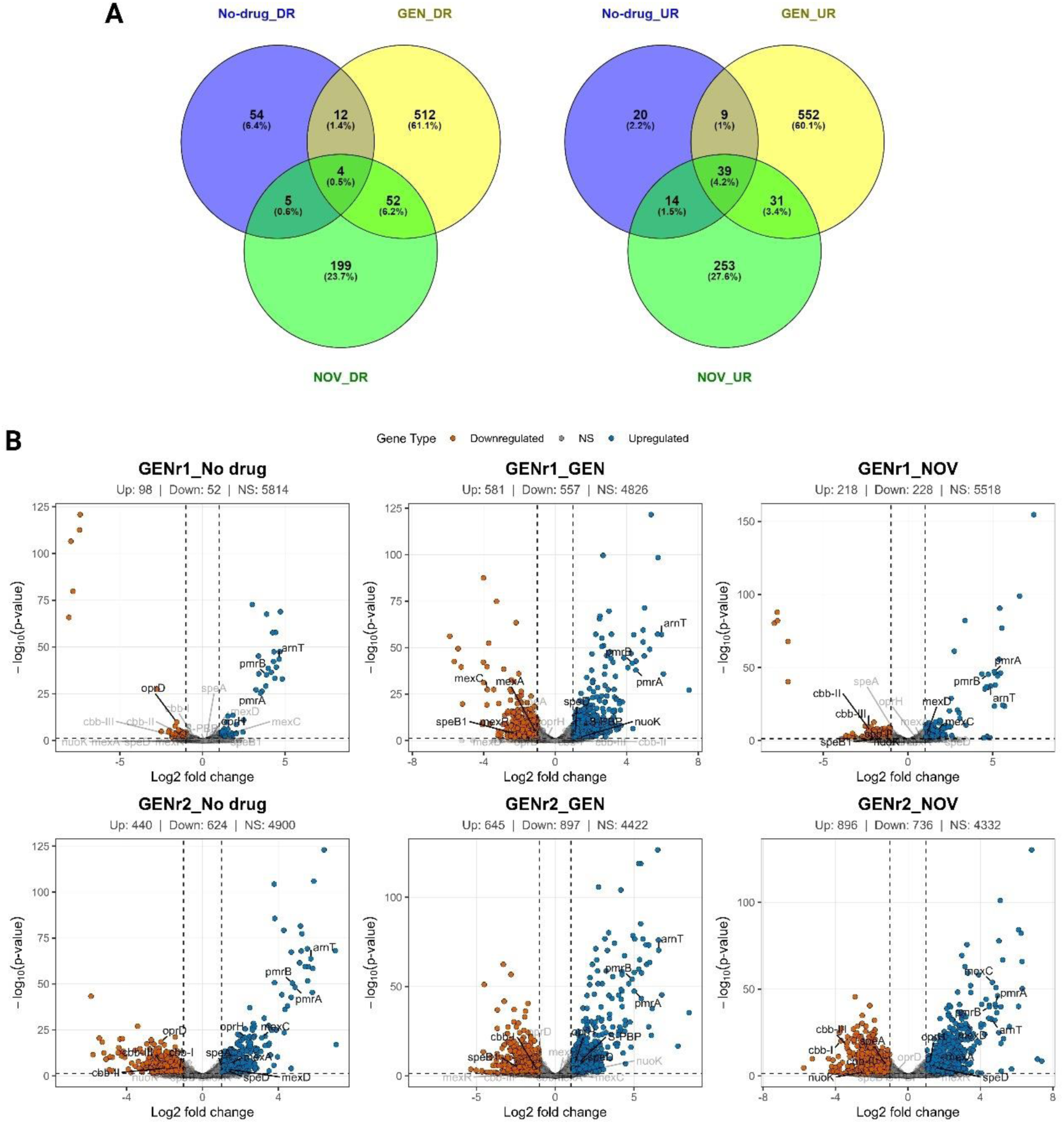
Transcriptomic analysis reveals mechanisms underlying GEN–NOV collateral sensitivity. **(a)** Venn diagrams showing the overlap of differentially expressed genes (DEGs) in GEN-adapted *P. aeruginosa* compared to wild-type PA14 under different conditions. **(b)** Volcano plots displaying DEGs in GEN-adapted bacteria relative to controls. Upregulated genes are shown in blue, downregulated genes in orange, and nonsignificant genes in gray. Numbers of significantly up- and downregulated genes are indicated above each plot.

Furthermore, we observed an upregulation in *pmrA, pmrB* and *arnT* genes in experimentally evolved GENr bacteria across all three conditions (Fig 5B). In general, PmrAB gain-of-function mutation and subsequent increased gene expression leads to the activation of arnBCADTEF operon, that adds L-Ara4N to lipid A, in turn neutralizes negative charges on the bacterial outer membrane (OM) (58). This reduces binding of cationic antimicrobials (e.g., polymyxins). Interestingly, earlier studies have shown using gene deletion experiments, the decreased susceptibility to aminoglycosides due to the *pmrAB* mutants was independent from operon arn, but rather depends on MexXY-OprM efflux and norspermidine production via the *speD2* and *speE2* gene cluster (58). Therefore, the observed GEN resistance mechanism maybe not be due to L-Ara4N lipid A modification. Instead, PmrAB must influence other targets that alter aminoglycoside susceptibility. Notably, we observed a *speD* (S-adenosyl-methionine decarboxylase) and (S-PBP) spermidine/putrescine-binding protein upregulation together with *speA* (arginine decarboxylase) and *speB1* gene downregulation in GENr following GEN treatment (Fig 5B). This represents a clear rewiring of polyamine flux with evidence for potential increased polyamine handling/transport, which is consistent with a polyamine coat/barrier that reduces aminoglycoside self-promoted uptake, exactly the arn-independent route noted earlier (58). Furthermore, gentamicin is a cationic aminoglycoside that needs to cross the OM through self-promoted uptake and (in part) porin-mediated routes. We found a downregulation in *oprD* gene expression that reduces the availability of a channel for GEN influx, additionally observed *oprH* gene upregulation in no treatment GENr cell which further stabilizes/insulates the OM, making it harder for GEN to breach the outer barrier (59, 60). Together, this porin remodeling means GEN has a harder time entering the GENr bacteria hence become resistant to GEN.

On the other hand, large lipophilic antibiotics such as novobiocin, macrolides or rifamycins can penetrate the outer membrane lipid bilayer via slow passive diffusion (61). Following NOV treatment, GENr upregulates RND efflux components (*mexA, mexC, mexD*), and *mexR* (repressor) is down under GEN treatment (Fig 5B) which declares that GERr cells are trying to pump. RND pumps such as MexAB-OprM and MexCD-OprJ function as proton/drug antiporters, they harness the energy from the PMF (62, 63). Without sufficient PMF, these pumps cannot drive drug export even if they are highly expressed (64). As PMF is low in GERr cells as detected earlier (Fig 4A), pumps underperform despite high mRNA, so NOV accumulates as a regulatory energetic mismatch. Furthermore, NOV treatment down regulates a broad range of electron transport/respiration genes (e.g., multiple nuo subunits; cbb3 type cytochrome c oxidase components), plus many transporters/porins (*oprG/E/L* down) (Fig 5B). Genes in *nuo-*operon and cbb3 type cytochrome c oxidase are integral part of energy metabolism (65, 66). OprG is the second smallest porin in PA and may allow transport of different hydrophobic molecules, additionally, OprE are found to be abundant in outer membrane vesicles of bacteria present in biofilm (67). This pattern may partially explain how NOV mediated collapsed respiratory flux leading even lower PMF and further reduces the permeability of the outer membrane. Further experiments are needed to validate this phenomenon. In general, this energetic weakness may further increase NOV activity through drug accumulation because of PMF-dependent efflux stalls.

Interestingly, under GEN, we observed some *nuo*/oxidative genes upregulation, which fits a compensatory attempt to rebuild PMF against a cationic antibiotic; yet the net phenotype still reads as low PMF in GENr cells. Earlier studies have also presented the role of PMF in CS between different antibiotics, including aminoglycoside antibiotics such as GEN. Reduced PMF across the bacterial membrane is strongly associated with aminoglycoside resistance in bacteria; however, it may reduce the activity of different PMF-dependent efflux pumps and leave the bacteria susceptible to other antibiotics (50). For example, compromised PMF due to the mutation in associated genes in bacteria may increase resistance to aminoglycoside antibiotics and may make them susceptible to beta-lactam antibiotics (68).

Overall, GEN resistance in PA emerges with an activated PmrAB program and envelope remodeling that collectively lowers PMF and reduces cationic aminoglycoside entry (Figs 6A and B). The same energetic state cripples PMF dependent RND efflux, driving intracellular accumulation of novobiocin and collateral hyper susceptibility (Figs 6A and C). Transcriptomics from GENr strains shows *pmrAB* upregulation across conditions, porin shifts (*oprH*↑, *oprD*↓), and NOV specific repression of respiratory modules (*nuo/cco*), while RND pumps are transcriptionally induced yet functionally constrained by low PMF. This PMF–efflux mismatch explains the observed GEN–NOV CS and predicts that sequence tuned therapy (GEN→NOV) at adequate NOV exposure may suppress emergence of GENr and relapse.

**Fig 6.**
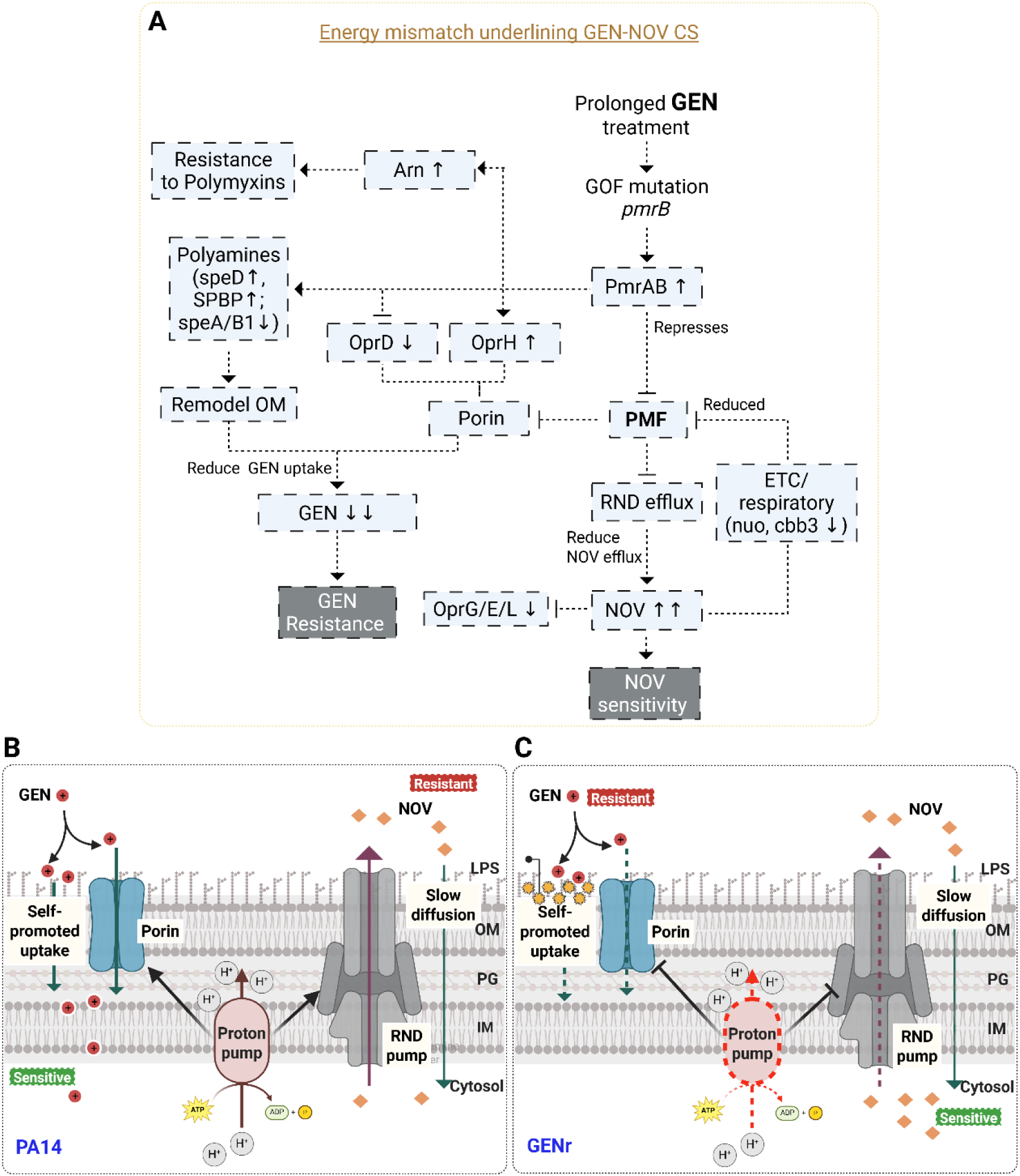
Transcriptomic and mechanistic model of GEN–NOV collateral sensitivity in *P. aeruginosa*. (A) Pathway model depicting how prolonged GEN exposure and gain-of-function (GOF) mutations in *pmrB* rewire gene expression and physiology in GEN-adapted *P. aeruginosa* (GENr). PmrAB upregulation activates polyamine pathways, remodels the outer membrane (OM), and shifts porin composition, collectively reducing GEN uptake. These changes, together with altered respiration and lowered proton motive force (PMF), impair energy-dependent resistance nodulation division (RND) efflux pumps. Thus, GENr cells display GEN resistance but NOV hypersusceptibility due to a regulatory–energetic mismatch. (B) Baseline PA14. In the wild-type strain, GEN efficiently enters via porins, and self-promoted uptake driven by PMF, while NOV diffuses slowly across the OM and is expelled by active RND efflux. This yields GEN sensitivity and NOV resistance. (C) GENr phenotype. In resistant derivatives, OM remodeling and reduced activity of porin decrease GEN entry, conferring resistance. Concurrently, compromised PMF diminishes RND efflux activity, causing NOV accumulation and hypersusceptibility. GOF; gain of function, ETC; electron transport chain, RND; resistance nodulation division efflux pump.

### GEN-NOV combination therapy to restrict the emergence of GEN resistance

The growing crisis of antibiotic resistance presents a significant hurdle in the effectiveness of antibiotic treatments. Antibiotic resistance is one of the major challenges in managing chronic infections, and it has been predicted that the number of deaths per year will rise to around 10 million in 2050 because of the rapid emergence of antibiotic resistance (69). As our study demonstrates NOV vulnerability in GENr PA, we wanted to see if the combination of NOV and GEN could reduce the emergence of GENr cells.

To this regard, we exposed the parental bacterial culture to both GEN and NOV. We administered NOV at suboptimal levels while gradually increasing the GEN concentration over time. Our experiment was structured into three distinct groups, such as, a no-drug control group, a group treated exclusively with GEN, and a final group receiving a combination of both NOV and GEN (Fig 7A). After 26 days of treatment, all three groups showed a similar count of viable cells. The control group exhibited the emergence of very negligible to low GENr cells. Compared to GEN alone, the NOV-GEN combination therapy showed a significant increase in GENr cells up to the 10-day mark. After 10 days, the number of GENr cells appeared comparable between these two treatment groups (Fig 7A). This outcome was quite unexpected and underscored the importance of careful consideration of any combined therapies. We theorize that the altered effect may appeared due to NOV’s target on the replication machinery (70). With NOV at suboptimal concentrations, the bacteria were not eradicated, but instead, experienced reduced efficacy in repairing DNA damage or mutations, leading to increased production of GENr cells.

**Fig 7.**
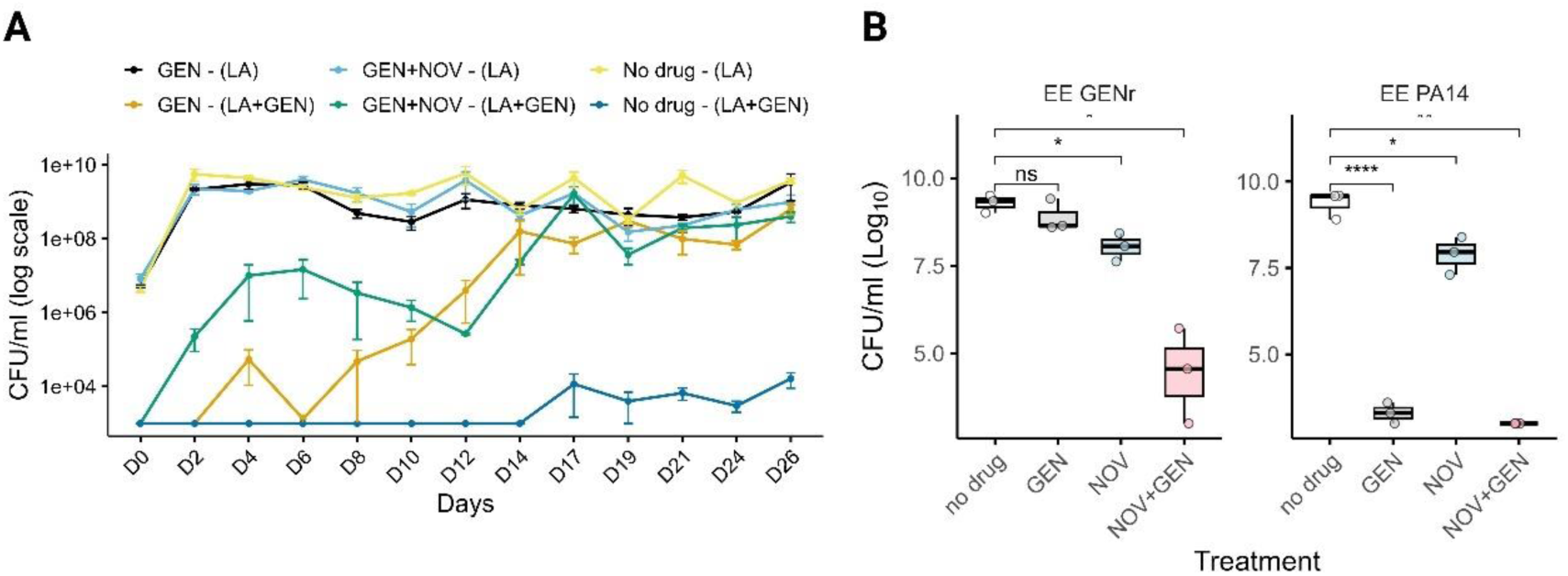
GEN–NOV combination therapy limits emergence of gentamicin-resistant cells. (A) Co-treatment assay showing total viable cells (LA plates) and GEN-resistant (GENr) populations (LA+GEN plates) under no-drug, GEN, or GEN+NOV conditions. (B) Sequential treatment assay in which GEN-adapted (EE GEN_r_) or vehicle control (EE PA14) cells were plated with GEN, NOV, or GEN+NOV. Data represent mean ± SD of three technical replicates. ‘****’ indicates *p* < 0.0001; ‘ns’ indicates not significant (two-tailed unpaired Student’s *t*-test).

Based on these data, we revised our treatment strategy. We increased the concentration of NOV and introduced sequential treatment instead of co-treatment. Along with no-drug control, first, parental bacteria were exposed to GEN alone, following the previously detailed experimental evolution approach. Then the evolved strains were treated with GEN alone, NOV alone, and GEN+NOV combined. Our findings showed no significant difference in the total number of viable cells between the two primary groups, GEN-adapted or no-drug control populations (Fig 7B). However, the GEN-adapted population exhibited a much higher number of GENr cells than the no-drug control group. Most notably, the combination of NOV and GEN resulted in a substantial reduction (*p*<0.0001) in GENr cells compared to no treatment control (Fig 6B). This suggests that a higher concentration of NOV, when used in combination with GEN, may effectively suppress the rise of GENr cells.

Combinatorial treatments are found to be effective in restricting resistance emergence and providing a better therapeutic treatment because of synergistic efficacy (71). Besides, considering the discovery of very few new antibiotics, combinatorial therapy offers potential treatment for infectious diseases using the existing antibiotics. The combinatorial treatment approach has been applied to manage different infectious diseases, such as HIV and tuberculosis, for many decades (38, 72, 73). Interestingly, in addition to reduced treatment cost and time, combinatorial treatment was found effective in restricting the emergence of antibiotic resistance by targeting collateral antibiotic sensitivity profiles (28, 74). For example, ciprofloxacin-adapted clinical isolates of PA were found to be sensitive to aztreonam, and the resistant mutant population was eliminated completely following the combinatorial treatment of ciprofloxacin-aztreonam (75). When CS is present, both concurrent and single day cycling treatments efficiently decrease resistance. This emphasizes the crucial role that drug administration sequence plays in determining the effectiveness of CS-based cycling therapies (76).

## Conclusion

Our data demonstrates that the acquisition of GEN resistance in PA leads to a reproducible CS to NOV. Using a combination of clinical isolates, experimentally evolved GENr strains, comparative genomics, and transcriptomic analyses, we show that GEN resistance is accompanied by *pmrB* gain-of-function mutation-associated outer membrane remodeling, reduced PMF, and impaired efflux activity. This energetic mismatch underlies the enhanced susceptibility of GENr bacteria to NOV. Importantly, sequential exposure to NOV following GEN adaptation effectively reduces GENr subpopulations, underscoring the therapeutic potential of exploiting CS relationships. Together, our results provide mechanistic insight into GEN–NOV CS and highlight the feasibility of leveraging evolutionary trade-offs to design rational antibiotic strategies. Incorporating CS-informed regimens may extend the lifespan of existing antibiotics and improve treatment outcomes for multidrug-resistant PA infections.

