## Supplemental figures 1 and 2 for "Proton motive force mediated efflux mismatch drives gentamycin-novobiocin collateral sensitivity in *Pseudomonas aeruginosa*"

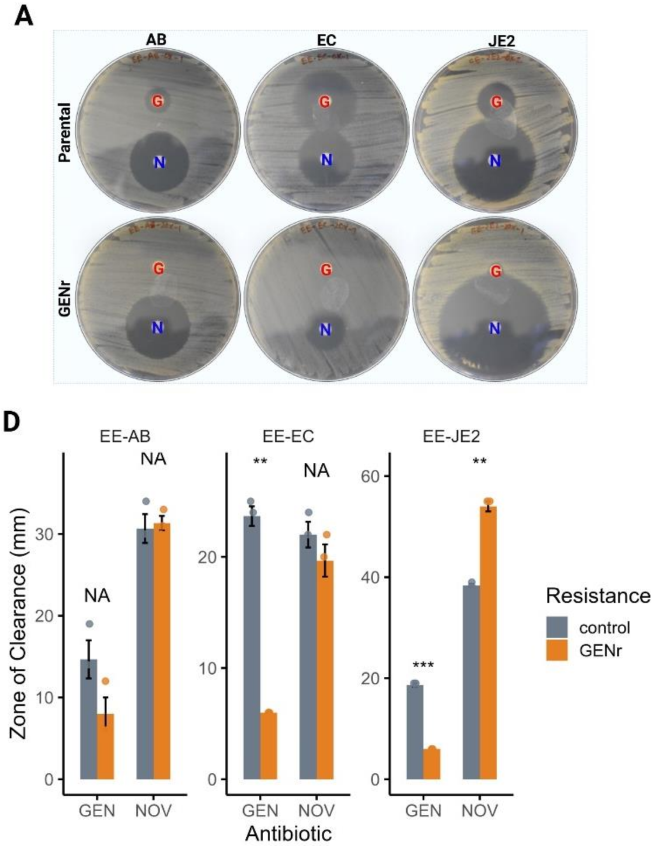


**Fig S1.** Sensitivity to novobiocin in GEN-resistant strains. (A) Disc diffusion assay of parental and GEN-adapted (GENr), *A. baumannii* (AB), *E. coli* (EC), and *S. aureus* (JE2) against GEN and NOV. (D) Quantification of clearance zones (mm) for GEN and NOV in experimental evolution (EE) strains compared with controls. Bars represent mean ± SD of three replicates. '**' indicates $p < 0.005$, '***' indicates $p < 0.0005$; NA = not applicable (two-tailed unpaired Student's t-test).

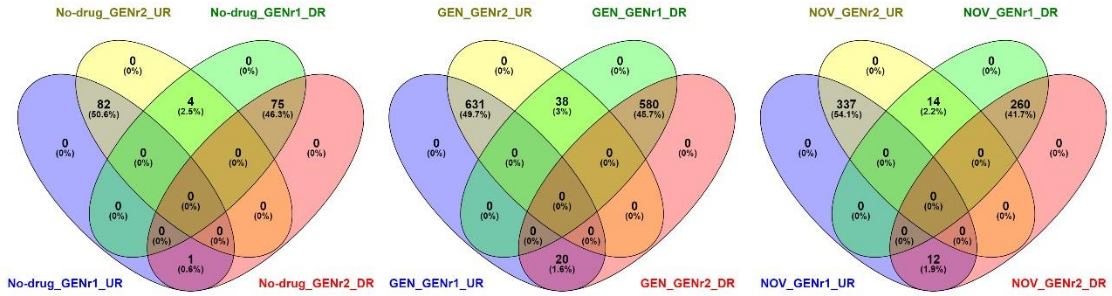


**Fig S2. Differential gene expression profiles in GENr strains under no-drug, GEN, and NOV treatment.** Venn diagrams show the overlap of upregulated (UR) and downregulated (DR) genes between two independently evolved GENr strains (GENr1, GENr2). (Left) No-drug control, (middle) GEN treatment, and (right) NOV treatment. The majority of DEGs are condition-specific, with limited overlap between strains or treatments.
